# Systems biology reveals NR2F6 and TGFB1 as key regulators of feed efficiency in beef cattle

**DOI:** 10.1101/360396

**Authors:** Pâmela A. Alexandre, Marina Naval-Sanchez, Laercio R. Porto-Neto, José Bento S. Ferraz, Antonio Reverter, Heidge Fukumasu

## Abstract

Systems biology approaches are used as strategy to uncover tissue-specific perturbations and regulatory genes related to complex phenotypes. We applied this approach to study feed efficiency (FE) in beef cattle, an important trait both economically and environmentally. Poly-A selected RNA of five tissues (adrenal gland, hypothalamus, liver, skeletal muscle and pituitary) of eighteen young bulls, selected for high and low FE, were sequenced (100bp, pared-end). From the 17,354 expressed genes, 1,317 were prioritized by five selection categories (differentially expressed, harbouring SNPs associated with FE, tissue-specific, secreted in plasma and key regulators) and used for network construction. NR2F6 and TGFB were identified and validated by motif discovery as key regulators of hepatic inflammatory response and muscle tissue development, respectively, two biological processes demonstrated to be associated to FE. Moreover, we indicated potential biomarkers of FE which are related to hormonal control of metabolism and sexual maturity. By using robust methodologies and validation strategies, we confirmed main biological processes related to FE in *Bos indicus* and indicated candidate genes as regulators or biomarkers of superior animals.

## Introduction

Since the domestication of the first species, animal selection aims to meet human needs and their changes over time. The current main selection goals in livestock production are increase of productivity, reduction of the environmental impact and reduction of competition for grains with human nutrition (Hayes *et al*, 2013). Thus, feed efficiency (FE) has become a relevant trait of study, as animals considered of high feed efficiency are those presenting reduced feed intake and lower production of methane and manure without compromising animal’s weight gain (Gerber *et al*, 2013). However, the incorporation of FE as selection criteria in animal breeding programs is costly and time consuming. Daily feed intake and weight gain for a large number of animals need to be recorded for at least 70 days to obtain accurate estimates of FE (Archer *et al*, 1997).

In the past years, several studies have been carried out with the aim to identify molecular markers associated with FE to enable the faster and cost-effectively identification of superior animals (de Oliveira *et al*, 2014; Rolf *et al*, 2011; Santana *et al*, 2014; Seabury *et al*, 2017). However, for each population, different biological processes seem to be identified (de Oliveira *et al*, 2014; Rolf *et al*, 2011; Santana *et al*, 2014; Seabury *et al*, 2017). Probably, that is because FE is a multifactorial trait and many different biological mechanisms seems to be involved in its regulation (Herd *et al*, 2004; Herd & Arthur, 2009). It has been indicated that high FE animals present increased mitochondrial function (Lancaster *et al*, 2014; Connor *et al*, 2010), less oxygen consumption (Gonano *et al*, 2014) and delayed puberty (Randel & Welsh, 2013; Shaffer *et al*, 2011; Fontoura *et al*, 2016). On the other hand, low FE animals have increased physical activity, ingestion frequency and stress (Francisco *et al*, 2015; Cafe *et al*, 2011; Kelly *et al*, 2010; Chen *et al*, 2014), increased leptin and cholesterol levels (Alexandre *et al*, 2015; Foote *et al*, 2016; Nkrumah *et al*, 2007; Mota *et al*, 2017), higher subcutaneous and visceral fat (Santana *et al*, 2012; Gomes *et al*, 2012; Mader *et al*, 2009), higher energy wastage as heat (Montanholi *et al*, 2010, 2009; Archer *et al*, 1999) and more hepatic lesions associated with inflammatory response (Alexandre *et al*, 2015; Paradis *et al*, 2015).

In the context of such a complex trait, we perform a multiple-tissue transcriptomic analyses of high and low FE Nellore cattle across tissues related to endocrine control of hanger/satiety, hydric and energy homeostasis, stress and immune response, physical and sexual activity, as is the case of hypothalamus-pituitary-adrenal axis and organs as liver and skeletal muscle. Based on gene co-expression across tissues and conditions we derived a regulatory network revealing NR2F6 and TGFB signalling as key regulators of hepatic inflammatory response and muscle tissue development, respectively. Next, we apply advanced motif discovery methods which i) validate that co-expressed genes are enriched for NR2F6 and TGFB signalling effector molecule SMAD3 binding sites in their 10KB upstream regions and ii) predict direct transcription factor (TF) – Target gene (TG) interactions at the sequence level. These binding interactions were experimentally validated with public TF ChIP-seq from ENCODE. Regulatory activity in the tissues of interest was also confirmed by performing an enrichment analysis on open chromatin tracks and histone chromatin marks across cell types and tissues in the human and cow genome. Moreover, we propose a hormonal control of differences in metabolism and sexual maturity between high and low FE animals, indicating potential biomarkers for further validation such as adrenomedullin, FSH, oxytocin, somatostatin and TSH.

## Results

### Multi-tissue transcriptomic data reveal differences between high and low feed efficient animals

Feed efficiency is a complex trait characterized by multiple distinct biological processes including metabolism, ingestion, digestion, physical activity and thermoregulation (Herd *et al*, 2004; Herd & Arthur, 2009). To study FE at transcriptional level we performed RNAseq of five tissues (i.e. adrenal gland, hypothalamus, liver, muscle and pituitary) from nine male bovines of high feed efficiency (HFE, characterized by low residual feed intake (RFI) (Koch *et al*, 1963)) and nine of low FE (LFE, characterized by high RFI). In total, we analysed 18 samples of liver, hypothalamus and pituitary; 17 of muscle and 15 of adrenal gland, yielding 13 million reads per sample on average (S1 Supporting Information). Gene expression was estimated for 24,616 genes present in the reference genome (UMD 3.1) and after quality control (refer to methods), 17,354 genes were identified as being expressed in at least one of the five tissues analysed.

Differential expression (DE) analysis between high and low FE animals resulted in 471 DE genes across tissues (P<0.001, S2 Supporting Information), namely, 111 in adrenal gland, 125 in hypothalamus, 91 in liver, 104 in muscle and 98 in pituitary (S3A-E Supporting Information). Although no significant functional enrichment was found for the 281 genes up-regulated in high feed efficiency, the 248 genes down-regulated presented a significant enrichment of GO terms such as response to hormone (Padj=5.43 x 10^-6^), regulation of hormone levels (Padj=3.48 x 10^-6^), cell communication (Padj=3.18 x 10^-4^), regulation of signaling receptor activity (Padj=3.20 x 10^-4^), hormone metabolic process (Padj=5.86 x 10^-4^), response to corticosteroid (Padj=6.28 x 10^-4^), regulation of secretion (Padj=7.2 x 10^-4^), response to lipopolysaccharide (Padj=7.9 x 10^-4^) and regulation of cell proliferation (Padj=1.86 x 10^-3^). Refer to S4 Supporting Information to see all enriched terms.

### Overlap between gene selection criteria prioritizes genes associated with feed efficiency

The genetic architecture behind complex traits involves a large variety of genes with coordinated expression pattern, which can be represented by gene regulatory networks as a blueprint to study their relationships and to identify central regulatory genes (Swami, 2009). Therefore, it is important to select relevant genes and gene families according to the phenotype of interest to be used for network analysis. We defined five categories of genes (see methods for further information) for inclusion in co-expression analysis: 1 - differentially expressed (DE), 2 - genes harbouring SNPs previously associated with FE (harbouring SNP), 3 - tissue specific (TS), 4 - genes coding proteins secreted in plasma by any of the five tissues analysed (secreted) and 5 - key regulators.

As reported before, we have identified 471 DE genes between high and low FE animals (Figure 1A, S5A Supporting Information). In addition, 267 genes were selected for harbouring SNPs previously associated with FE, as not only differences in expression levels can influence the phenotype but also polymorphism in the DNA sequence that can alter the translated protein behaviour (S5B Supporting Information). Moreover, 396 were selected for being tissue specific (refer to methods for definition); 22 in adrenal gland, 32 in hypothalamus, 215 in liver, 218 in muscle and 9 in pituitary (S5C Supporting Information). A total of 244 genes coding proteins secreted in plasma were selected because of their potential as biomarkers of FE (S5D Supporting Information). From those, 135 had liver as the tissue of maximum expression and were functionally enriched for GO terms such as complement activation (Padj=1.82 x 10^-19^), regulation of acute inflammatory response (Padj=1.89 x 10^-14^), innate immune response (Padj=9.71 x 10^-12^), negative regulation of endopeptidase activity (Padj=2.35 x 10^-10^), platelet degranulation (Padj=1.08 x 10^-10^), regulation of coagulation (Padj=3.39 x 10^-9^), triglyceride homeostasis (Padj=1.23 x 10^-6^), cholesterol efflux (Padj=1.03E^-5^) (S6 Supporting Information). Finally, from 1570 potential regulators in public available TFdb, 78 were identified as key regulators of the genes selected by all the other categories, i.e. 78 genes presented a coordinated expression level with many of the genes in the network reflecting a tight control of expression pattern across tissues (S5E Supporting Information).

**Figure 1.**
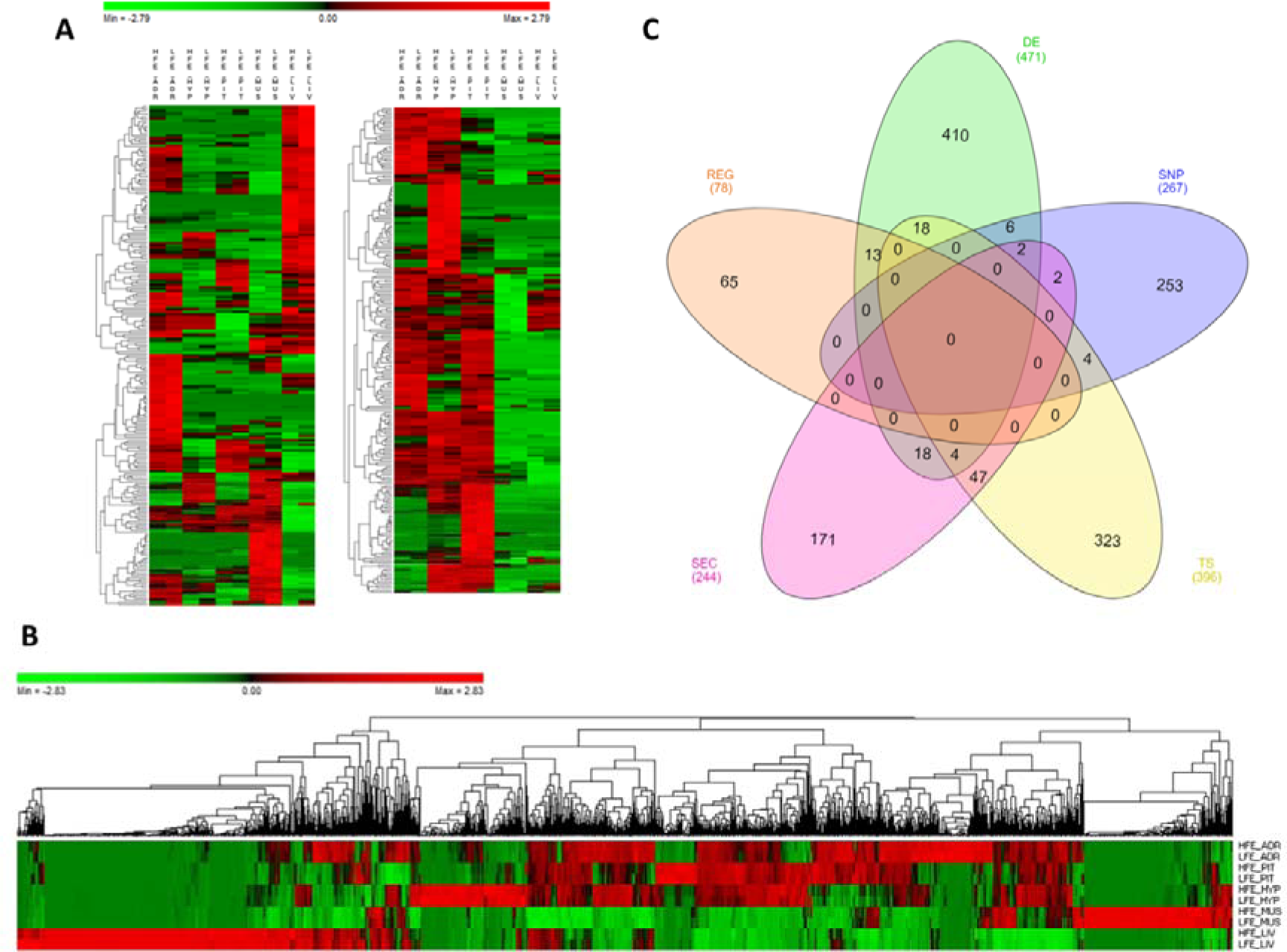
Genes selected for co-expression network construction. **A)** Heatmap of normalized mean expression (NME) of 471 differentially expressed (DE) genes between high (HFE) and low (LFE) feed efficient animals in adrenal gland (ADR), hypothalamus (HYP), liver (LIV), muscle (MUS) and pituitary (PIT). Genes (rows) and samples (columns) are organized by hierarchical clustering based on Euclidean distances. **B)** NME heatmap of all 1,317 genes selected for network construction. Genes (columns) and samples (rows) are organized by hierarchical clustering based on Euclidean distances. **C)** Venn diagram of 1,317 genes selected for network construction. The inclusion criteria for selecting genes were divided in five categories: differentially expressed genes (DE), tissue specific genes (TS), genes harbouring SNPs reported by literature as being associated with feed efficiency in beef cattle (SNP), genes encoding proteins secreted by at least one of the tissues in plasma (SEC) and key regulators (REG). Numbers between brackets indicate the total number of genes in each category.

Considering all the inclusion criteria, 1,317 genes were selected to be included in co-expression network analysis (Figure 1B, S7 Supporting Information), some of them selected by more than one category (Figure 1C). Regarding DE genes, six of them were also reported before as harbouring SNPs associated with the phenotype (*LUZP2, MAOB, SFRS5, SLC24A2, SOCS3* and *WIF1*) and 13 of them were key regulators (*HOPX, PITX1, CRYM, PLCD1, ND6, cytb, ND1, MT-ND4L, ND5, ATP8, ND4, ENSBTAG00000046711* and *ENSBTAG00000048135*). Many of the genes that are both DE and regulators are involved in respiratory chain (*ND6, cytb, ND1, MT-ND4L, ND5, ATP8* and *ND4*) and were all up-regulated in high FE group.

Considering both DE and secreted genes, 18 were identified (*NOV, SPP1, CTGF, OXT*, *PTX3, VGF, CCL21, COL1A2, PGF, SOD3, SERPINE1, PRL*, *PON1, SST, JCHAIN, PCOLCE, IGFBP6* and *SCG2*). In addition, four genes were DE, secreted and tissue specific, two from liver (*CXCL3* and *IGFBP1*) and two from pituitary (*NPY* and *CYP17A1*). Genes *RARRES2* and *PENK* (proenkephalin) were DE, secreted and had been previously reported as harbouring SNP associated with FE [30, AnimalQTLdb]. Other DE genes worthy to highlight, due to their well-known role in metabolic processes, are *AMH* (anti-mullerian hormone), *TSHB* (thyroid stimulating hormone beta), *FGF21* (Fibroblast growth factor 21) and *FST* (follistatin), up-regulated in high FE group, and *PMCH* (pro-melanin concentrating hormone), *ADM* (adrenomedullin) and *FSHB* (follicle stimulating hormone beta), up-regulated in low FE group.

### Co-expression network reveals regulatory genes and biological processes related to feed efficiency

The co-expression network (Figure 2) was composed by 1,317 genes and 91,932 connections, with a mean of 70 connections per gene. Most of the connections (51%) involved a DE gene and 23% of those were between two DE genes. Tissue specific (TS) gene were involved in 49% of the connections with 119 connections per gene in average, which was higher than the overall network mean and reflects the close relationship between genes involved in tissue specific functions. Key regulators was the least represented category in the network (only 78 genes) but accounted for 11% of the connections in the network with the highest value of mean connections per gene, 131 connections, which is in accordance with their regulatory role. Regarding the connections within tissues, when we ranked all the genes in the network by the number of connections and looked at the top 50 genes, 29 were from liver, 15 were from muscle and 3, 2 and 1 were from pituitary, adrenal gland and hypothalamus, respectively. This result indicates a very well-coordinated expression pattern in liver and muscle that could be a reflex of the number of TS genes in those tissues and the presence of central regulatory genes coordinating the expression of many other genes.

**Figure 2.**
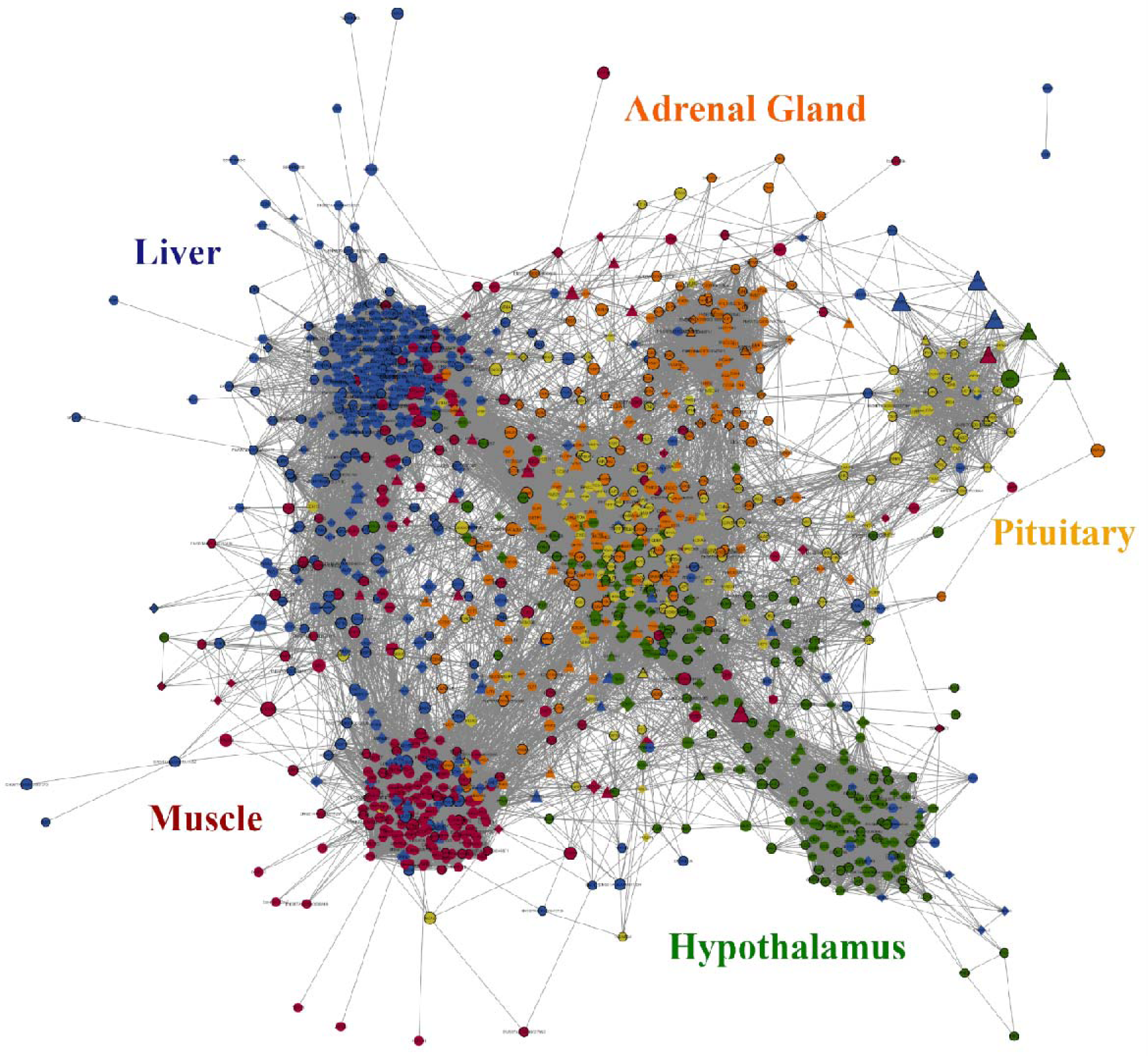
Gene co-expression network constructed using PCIT algorithm on 1,317 selected genes (see methods). Nodes with diamond shape correspond to secreted proteins coding genes and triangles correspond to key regulators; all the other genes are represented by ellipses. Nodes with black borders are differentially expressed between high and low feed efficiency. Colours are relative to the tissue of maximum expression: blue represent liver, red represent muscle, yellow represent pituitary, green represent hypothalamus and orange represent adrenal gland. The size of the nodules is relative to the normalized mean expression values in all samples. Only correlations above 0.9 and bellow −0.9 and its respective genes are shown in this figure.

In the network (Figure 2), genes grouped together by tissue which was mostly driven by TS genes. As mentioned before, most of the secreted proteins coding genes were locate in the liver. Most of the key regulators were located peripherally in relation to the clusters which could be reflecting their regulatory nature independent of tissue specificity. Despite that, some regulators draw attention because of their high number of connections.

The top five most connected regulators were *EPC1, NR2F6, MED21, ENSBTAG00000031687* and *CTBP1*, varying from 317 to 284 connections. They were all first neighbours of each other and were connected mainly to genes with higher expression in liver and essentially enriched for acute inflammatory response (Padj=4.5 x 10^-13^, S8 Supporting Information). The next most connected regulator is *TGFB1* with 217 connections. It is mainly connected to genes from muscle that are primarily enriched for muscle organ development (Padj=6.87 x 10^-5^) and striated muscle contraction (Padj=1.39 x 10^-5^, S9 Supporting Information). Besides indicating main regulator genes, network approach can be useful to access the role of specific genes. For instance, gene *FGF21*, a hormone up regulated in liver of high FE animals, is directly connected to genes enriched for plasma lipoprotein particle remodelling, regulation of lipoprotein oxidation and cholesterol efflux (Padj=5.64E-3, S10 Supporting Information). Indeed, according to the literature, this gene is associated to decrease in body weight, blood triglycerides and LDL-cholesterol (Cheung & Deng, 2014).

### Motif discovery confirms NR2F6 as a key regulator of liver transcriptional changes between high and low feed efficiency

By means of the power-law theory, co-expression networks present many nodules with few connections and few central nodules with many connections (de la Fuente, 2010), being the last ones indicated as central regulatory genes responsible for the transcriptional changes between the divergent phenotypes analysed. In our study, the most connected regulators were indicated, together with their target genes, i.e. their first neighbours in the network. Those genes are a mixture of direct and indirect regulator targets. In order to validate the regulatory role of the most connected regulators in the network and identify their core direct targets we performed motif discovery in their co-expressed target genes. It is noteworthy motif discovery should confirm the presence of DNA motifs of a TF in the regulatory regions of co-expressed genes. From the top five most connected regulators from our previous co-expression analysis, only NR2F6 has the ability to bind DNA. In contrast, the other four regulators act mainly as cofactors (corepressor, i.e. CTBP1; coactivator, i.e. MED21; or histones modifier, i.e. EPC1), that is co-binding through protein-protein interactions.

The analysis of 313 co-expressed genes with NR2F6 yield the Nuclear Factor motif HNF4-NR2F2 (transfac_pro-M01031) as the second motif most enriched out of 9732 PWMs with a Normalized Enrichment Score (NES) of 7.99 (Figure 3B). In addition, a total of 19 motifs associated with HNF4-NR2F2 were enriched in the dataset, associating HNF4-NR2F2 to 168 direct target genes (Figure 3C). Due to motif redundancy or highly similarity between a plethora of TFs, these motifs can be associated with multiple TFs from HNF4 (direct) to several nuclear factors such as NR2F6 (motif similarity score FDR 1.414E-5). However, our co-expression analysis strongly indicates NR2F6 is the key TF, since it was the TF with the highest number of nodes in the co-expression network (Figure 3C) and neither HNF4 nor NR2F2 were prioritized by any selection category to be included in the network.

**Figure 3.**
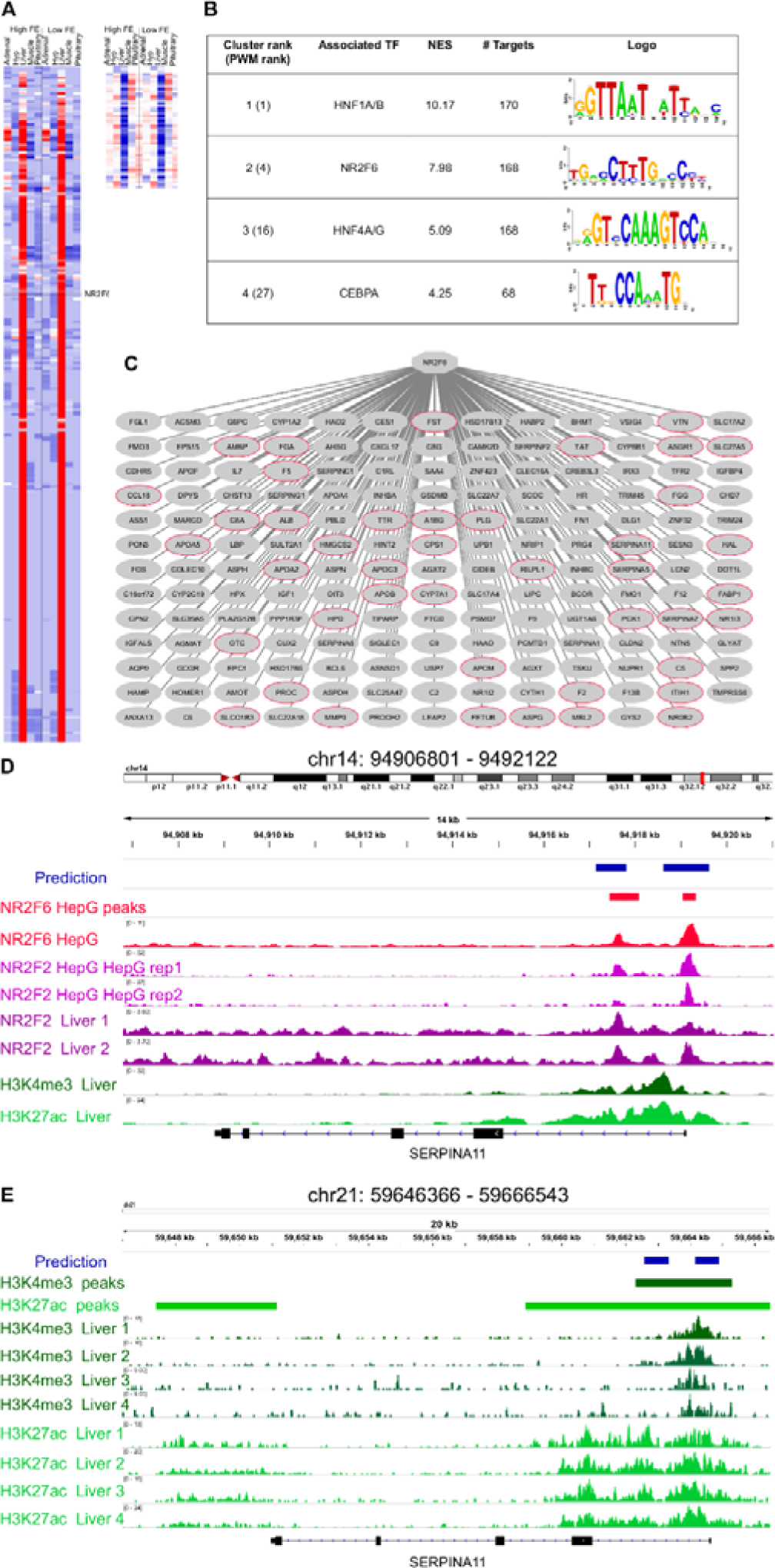
Mapping of NR2F6 direct targets. A) Heatmap of the 313 genes coexpressed with NR2F6 across all samples (derived from the co-expression analysis), B) i-cisTarget motif discovery results on the genes shown in (A), C) Predicted NR2F6 targetome. A red node indicates genes known to be targeted my NR2F6 in human Hepatocytes. D) Example of predicted NR2F6 target regions for SERPINA1 gene. The predicted enhancer overlaps the exact position for NR2F6 and NR2F2 binding in HepG sites from ENCODE dataset as well as histone chromatin marks related with active regulatory regions, namely H3K27ac, and promoters, H3K4me3 in human primary tissue from RoadMap Epigenetics E) The enhancer prediction in cow coordinates (bosTau6) overlaps a region marked with H3K4me3 in cow liver (Villar *et al*, 2015).

Each of the NR2F6 inferred direct target genes contain one or more predicted enhancers, i.e. regions with high-scoring motif binding sites for NR2F6 or TFs with highly similar motifs. To validate the binding of these genomic regions by NR2F6 or TFs with highly similar motifs to NR2F6 we performed a region enrichment analysis of our predicted NR2F6 binding sequences against public TF ChiP-seq bound regions in human cell lines from ENCODE the ENCODE consortium (1394 TF binding site tracks). This analysis, confirms the experimental binding of TFs with similar binding as NR2F6 in HepG2 cells, HNF4A (NES=8.57), HNF4G (NES=7.83), RXRA (NES=6.85), and NR2F2 (NES=4.45) as the most enriched tracks (S11 Supporting Information). Recent NR2F6 ChIP-seq data in HepG also confirms an enrichment for NR2F6 (Figure 3D), indicating predicted NR2F6 binding regions are experimentally bound by NR2F6 in hepatocyte cell lines (Figure 3D).

Next, to validate that the NR2F6 binding in those regions is functional in liver we performed an enrichment analysis for open-chromatin (tracks=655) and histone modifications (tracks =2450) related to active regulatory elements (S12 Supporting Information). This analysis yielded FAIRE-seq on HepG2 cell lines and H3K9ac and H3K4me3 in adult liver (E066 Roadmap Epigenomics Track) as the most enriched tracks respectively, strongly indicating not only predicted target enhancers are bound by NR2F6 in Hepatocyte cell lines but these regulatory regions are functionally active in hepatocytes and human liver (Figure 3D).

Regarding the cow genome, a recent open-chromatin study (Villar *et al*, 2015) has delineated the map active promoters and enhancers by H3K4me3 and H3K27ac ChIP-seq in cow liver resulting in 13796 promoter and 45786 enhancers (S13 Supporting Information). We performed an enrichment analysis of predicted NR2F6 enhancers converted to cow coordinates (n=779) resulting in 446 regions being identified as functional regulatory regions in cow liver. This number is significantly higher compared to only 43 regions are expected to overlap by random (1000 permutation tests) (Figure 3E).

Finally, in addition to NR2F6 motif, HNF1A motif was found as a potential co-regulator in liver, in particular swissregulon-HNF1A.p2 with a NES =10.17 and in total 20 enriched motifs and 170 direct targets were associated to HNF1A (Figure 3B). HNF1 is a master regulator of liver gene expression (Tronche & Yaniv, 1992), thus making its finding justified.

### Motif discovery validates TGF-beta signalling through Smad3/MyoD1 binding as drivers of transcriptional differences in muscle of divergent feed efficient cattle

The analysis of the 217 genes co-expressed with *TGFB1* (Figure 4A) showed most target genes motifs were enriched for master regulators of muscle differentiation, namely, *MEF2* (NES=10.42) a MADS box Transcription factor with 148 target genes, and *MYOD1* (NES=8.12), a bHLH transcription factor (CANNTG) with 135 direct target genes (Figure 4B, S14 Supporting Information). To evaluate the precision of our predicted *MYOD1* (bHLH) target genes we assessed how many of these TF-TG relationships had been previously experimentally reported. Based on *MYOD1* ChIP-seq binding in mouse myotubules, 86 genes had already been associated with *MYOD1* resulting in 63% success rate (hypergeometric test 1.72E-22). *SMAD3*, the effector molecule of *TGFB1* signalling is known to recruit *MYOD1* to drive transcriptional changes during muscle differentiation (Mullen *et al*, 2011). Thus, we evaluated whether predicted *MYOD1* target genes were enriched for known *SMAD3* target genes resulting in 21 out of 135 *MYOD1* predicted target genes presented *SMAD3* ChIP-seq binding in myotubes. Thus indicating there is a statistically significant association between *MYOD1* target genes and *SMAD3* target genes in myotubes (hypergeometric test 1.98 E-6) (Figure 4C) (Mullen *et al*, 2011) In contrast, no significant association was found between predicted *MYOD1* target genes in this study and *SMAD3* target genes in other cell lines, such as pro-B and ES cell (hypergeometric test 0.056 and 0.076, respectively) (Mullen *et al*, 2011). That is in agreement that the effect of TGFB signalling driven by SMAD3 DNA binding is tissue-specific (Liu *et al*, 2001). Our analysis predicted 621 potential *MYOD1* binding sites, of which 114 (18%) and 153 (24.5%) present a *MYOD1* ChIP-seq signal in mouse C2C12 myotubes cells (Mullen *et al*, 2011) and in primary myotubes (Cao *et al*, 2009), respectively.

**Figure 4.**
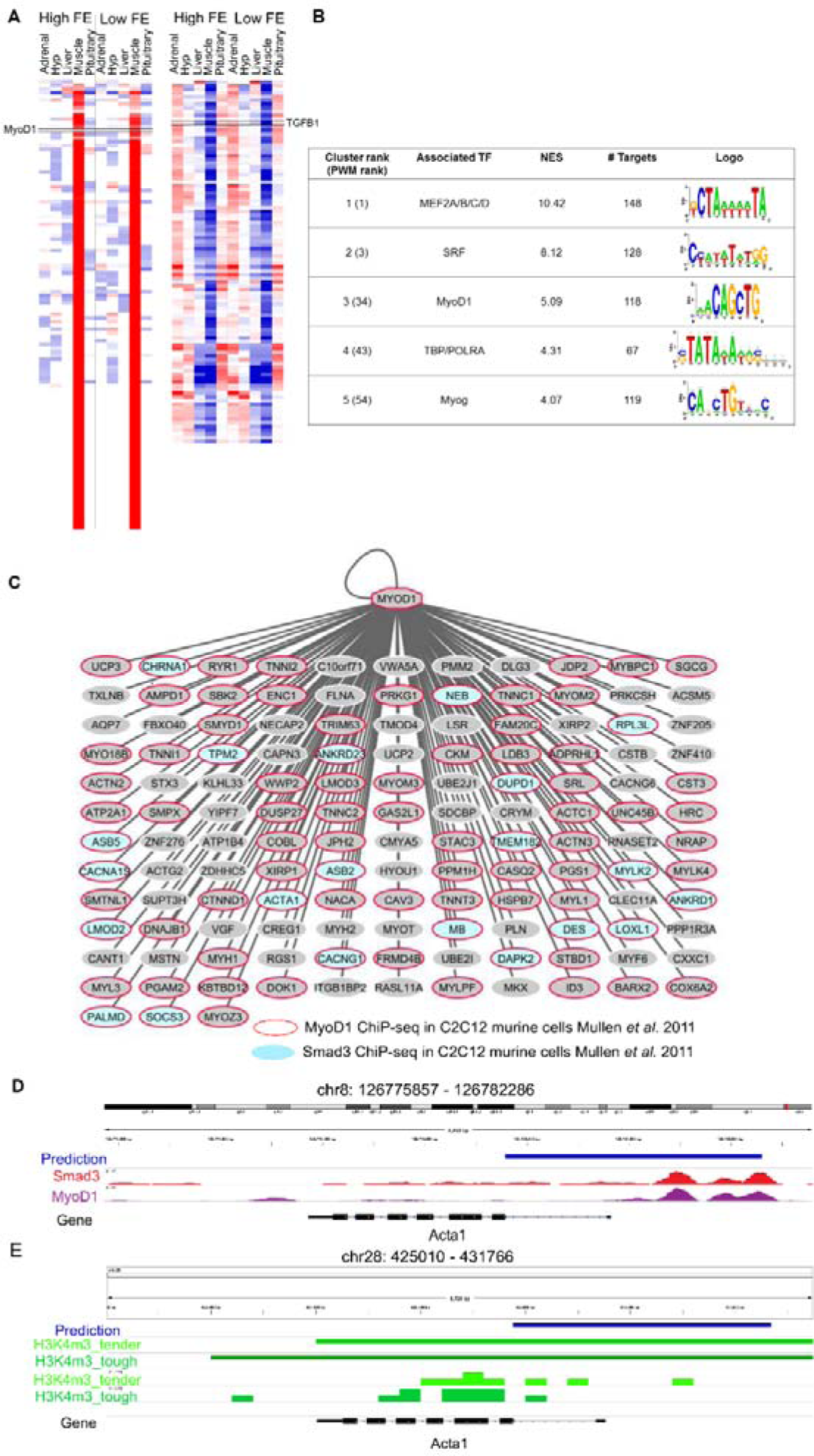
Mapping the downstream network of TGFB signaling through SMAD3/MyoD1 DNA binding. A) Heatmap of the 217 genes coexpressed with TGFB1 (derived from the co-expression analysis). B) i-cisTarget motif discovery results on the genes shown in (A), C) Predicted MyoD targetome. A red node indicate genes know to be targeted my MyoD1 in murine myotubes (Mullen *et al*, 2011). Blue nodes indicate genes to be targeted by SMAD3, the effector DNA binding molecular of TGFB signalling, in murine myotubes (Mullen *et al*, 2011). D) Example of predicted MyoD1 target regions for Acta1 gene. The predicted enhancer overlaps the exact position for SMAD3 and MyoD1 ChIP-seq binding in murine myotubes (Mullen *et al*, 2011). E) The enhancer prediction in cow coordinates (bosTau6) overlaps a promoter region marked with H3K4me3 in muscle tissue in cow (Cao *et al*, 2009).

Finally, we evaluate whether predicted MYOD1 binding regions were regulatory regions active in muscle cells across different species, namely human (S15 Supporting Information), mouse (S16 and S17 Supporting Information) and cow (S18 Supporting Information). To tackle this issue we performed an enrichment analysis across 2113 open-chromatin ENCODE tracks. This analysis resulted in a clear enrichment of our predicted MYOD1 binding regions with H3K27ac (NES=15.98) and H3K9ac (NES=8.78) regions in skeletal muscle (Figure 4D). Both chromatin marks are associated with active transcription, H3K27ac related to active enhancers and H3K9ac related to active gene transcription (Shin *et al*, 2012) thus, validating most of our enhancer predictions in human are active in skeletal muscle. Once in cow, we assess the overlap of predicted MYOD1 enhancers and promoter regions in cow muscle experimentally detected with H3K4me3 (Cao *et al*, 2009). This resulted in 282 regions out of 671 (42 %) overlap when only 11 regions are expected to overlap by random 1000 permutation test) (Figure 4E).

Finally, we evaluate whether predicted MYOD1 binding regions were regulatory regions active in muscle cells across different species, namely human (S15 Supporting Information), mouse (S16 and S17 Supporting Information) and cow (S18 Supporting Information). To tackle this issue we performed an enrichment analysis across 2113 open-chromatin ENCODE tracks. This analysis resulted in a clear enrichment of our predicted MYOD1 binding regions with H3K27ac (NES=15.98) and H3K9ac (NES=8.78) regions in skeletal muscle (Figure 4D). Both chromatin marks are associated with active transcription, H3K27ac related to active enhancers and H3K9ac related to active gene transcription (Shin *et al*, 2012) thus, validating most of our enhancer predictions in human are active in skeletal muscle. Once in cow, we assess the overlap of predicted MYOD1 enhancers and promoter regions in cow muscle experimentally detected with H3K4me3 (Cao *et al*, 2009). This resulted in 282 regions out of 671 (42 %) overlap when only 11 regions are expected to overlap by random 1000 permutation test) (Figure 4E).

## Differential co-expression

Although the general co-expression network give us important insights about regulatory genes and their behaviour, by creating specific networks for high and low FE and comparing the connectivity of the genes in each one, we can identify genes that change their behaviour depending on the situation, moving from highly connected to lowly connected and vice-versa. We were able to identify 87 differentially connected genes between high and low FE (P<0.05); 63 mainly expressed in liver, 19 in muscle and 3, 1 and 1 in hypothalamus, adrenal gland and pituitary, respectively (S19 Supporting Information). Those genes were enriched for terms such as regulation of blood coagulation (Padj=3.14 x 10^-10^), fibrinolysis (Padj=7.71 x10^-7^), platelet degranulation (Padj=7.49 x10^-6^), regulation of peptidase activity (Padj=6.16 x 10^-4^), antimicrobial humoral response (Padj=2.49 x10^-3^), acute inflammatory response (Padj=2.18 x10^-4^) and induction of bacterial agglutination (Padj=3.58 x 10^-2^) (S20 Supporting Information). It is important to highlight 20 of the differentially connected genes were also differentially expressed (Table 1) and three of them, i.e. *SST, JCHAIN* and *IGFBP1*, were secreted in plasma as well, which make them very promising potential biomarkers.

**Table 1.** Differentially connected and differentially expressed genes between high and low feed efficiency.

| Gene name | Number of connections |  | Category* | Tissue of maximum expression | Tissue of differential expression |
| --- | --- | --- | --- | --- | --- |
|  | Low feed efficiency | High feed efficiency |  |  |  |
| SST | 0 | 45 | DE, SEC | Hypothalamus | Hypothalamus |
| SNORA73 | 41 | 108 | DE | Liver | Liver |
| ENSBTAG00000047700 | 56 | 111 | DE | Liver | Liver |
| ENSBTAG00000047121 | 62 | 111 | DE | Liver | Liver |
| ENSBTAG00000047816 | 53 | 96 | DE | Liver | Liver |
| ENSBTAG00000039928 | 50 | 89 | DE | Liver | Liver |
| ANXA13 | 115 | 63 | DE | Liver | Liver |
| FST | 113 | 56 | DE | Liver | Liver |
| PBLD | 115 | 55 | DE | Liver | Liver |
| ENSBTAG00000021368 | 95 | 0 | DE | Liver | Liver |
| JCHAIN | 52 | 113 | DE, SEC | Liver | Liver |
| IGFBP1 | 55 | 0 | DE, TS, SEC | Liver | Liver |
| SBK2 | 0 | 70 | DE | Muscle | Muscle |
| ACTC1 | 54 | 0 | DE | Muscle | Muscle |
| MYH1 | 0 | 47 | DE, TS | Muscle | Muscle |
| HR | 119 | 50 | DE | Pituitary | Muscle |
| TAGLN | 83 | 31 | DE | Adrenal | Muscle, Pituitary |
| SFRP2 | 41 | 91 | DE | Hypothalamus | Pituitary |
| FN1 | 119 | 69 | DE | Liver | Pituitary |
| CAV1 | 98 | 50 | DE | Muscle | Pituitary |

\*Differentially expresses genes between high and low feed efficiency (DE), tissue specific genes (TS) and genes encoding proteins secreted in plasma (SEC).

Comparing the two networks there was no large difference regarding the number of genes and connections. While high FE network contained 1,074 genes in total and 28,018 connections, low FE network was composed by 1,098 genes and 30,705 connections. For all tissues, low FE networks showed more connections but the difference is slight, being the bigger difference of 44 versus 40 connections per gene in liver.

## Discussion

Feed efficiency is a complex trait, regulated by several biological processes. Thus, the indication of genomic regions associated with this phenotype, as well as regulators genes and biomarkers to select superior animals and to direct management decisions is still a great challenge. In this work, multi-tissue transcriptomic data of high and low feed efficient Nellore bulls were analysed through robust co-expression network methodologies in order to uncover some of the biology that governs this traits and put forward candidate genes to be focus of further research. In this sense, the validation of target genes of main transcription factors (key regulators) in the network by motif search proves the efficacy of the methodology for network construction and prioritizes some transcription factors as central regulators (Aerts *et al*, 2010; Naval-Sańchez *et al*, 2013; Potier *et al*, 2014). Moreover, the addition of a category of genes coding proteins secreted in plasma in the co-expression analysis highlight genes with potential to be explored as biomarkers of feed efficiency. We were able to identify genes related to main biological process associated with feed efficiency and indicate key regulators, as can be seen on the following lines.

Firstly, it is worthy to mention the 98 animals used to select the high and low FE groups in this study have been previously analysed regarding several phenotypic and molecular measures (Alexandre *et al*, 2015; Mota *et al*, 2017; Novais *et al*, 2018). It was observed high and low FE groups had similar body weight gain, carcass yield and loin eye area but low FE animals had higher feed intake, greater fat deposition, higher serum cholesterol levels, as well as hepatic inflammatory response, indicated by transcriptome analysis of liver biopsy and proved by the higher number of periportal mononuclear infiltrate (histopathology) and increased serum gamma-glutamyl-transferase (GGT, a biomarker of liver injury) in this group (Alexandre *et al*, 2015). In the present study, the simultaneous analysis of five distinct tissues revealed a prominence of the hepatic tissue. Liver presented the most connected genes in the network, the higher number of differentially connected genes and higher number of secreted genes, which although can be explained by its biological function, are enriched mostly for terms related to lipid homeostasis and inflammatory response. Moreover, the top five most connected regulators in the network are co-expressed mainly with genes highly expressed in liver and also enriched for inflammatory response.

The relationship between FE and genes or pathways related to immune response and lipid metabolism is becoming more evident, as recent studies also reported it in beef cattle (Karisa *et al*, 2014; Weber *et al*, 2016; Paradis *et al*, 2015; Zarek *et al*, 2017; Mukiibi *et al*, 2018) and pigs (Gondret *et al*, 2017; Ramayo-Caldas *et al*, 2018). In our previous work (Alexandre *et al*, 2015), we proposed increased liver lesions associated with higher inflammatory response in liver of low FE animals could be due to increased lipogenesis and/or higher bacterial infection in the liver. While further evidence is needed to test these hypotheses, the enrichment of terms such as induction of bacterial agglutination and response to lipopolysaccharide makes bacterial infection a strong possibility. Indeed, pigs with low FE were reported to have higher intestinal inflammation, neutrophil infiltration biomarkers and increased serum endotoxin (lipopolysaccharide and other bacterial products) which could be related to increased bacterial infection or to decreased capacity to neutralize endotoxins (Mani *et al*, 2013). The authors hypothesized differences in bacterial population could partially explain the increase in circulating endotoxins, which could also be true for cattle given that differences in intestinal and ruminal bacterial population between high and low FE animals have already been reported (Myer *et al*, 2015, 2016). Furthermore, the literature reports lipopolysaccharides (LPS) may cause up-regulation of adrenomedullin (*ADM*) hormone (Shindo *et al*, 1998), an up-regulated gene in low FE individuals as showed here. It was also demonstrated in rats that intravenous infusion of LPS caused up-regulation of *ADM* in ileum, liver, lung, aorta, skeletal muscle and blood vessels (Shoji *et al*, 1995) whereas in our study, *ADM* presented differential expression in muscle, but not in liver.

Against pathogen invasion, a tightly regulated adaptive immune response must be triggered in order to allow T lymphocytes to produce cytokines or chemokines and B cells to differentiate and produce antibodies (Hermann-Kleiter & Baier, 2014). This regulation is known to be strongly influenced by the expression level and transcriptional activity of several nuclear receptors, including the NR2F-family, which consists of three orphan receptors: NR2F1, NR2F2 and NR2F6 (Hermann-Kleiter & Baier, 2014). Those receptors present highly conserved DNA and ligand binding domains among each other and across species (Pereira *et al*, 2000), and all three are expressed in adaptive and immune cells (Hermann-Kleiter & Baier, 2014). In our study, NR2F6 appeared as the second most connected regulator gene in the network while the other family members, although present in our expression data, were not selected by any of our inclusion criteria, thus indicating they might not be so relevant in our conditions. Indeed, NR2F6 appears to be a critical regulatory factor in the adaptive immune system by directly repressing the transcription of key cytokine genes in T effector cells (Hermann-Kleiter *et al*, 2008; Klepsch *et al*, 2016). The role of NR2F6 as a key regulator of inflammatory response in our network was validated at gene level by the identification of the binding motif HNF4-NR2F2 (transfac_pro-M01031) as one of the most enriched in NR2F6 target genes, due to the high similarity between NR2F2 and NR2F6 biding sites. Furthermore, using open chromatin data public available, we provided experimental evidence of the binding of TFs with highly similar binding motifs as NR2F6 in hepatocyte cells in humans and in cattle, thus, indicating predicted target enhancers are functional in this tissue.

Another regulator prioritized in our analysis is *TGFB1*, the sixth most connected gene in the co-expression network, and a potential driver of transcriptional changes between high and low FE cattle in muscle. This gene has been previously pointed as a master regulator of FE in beef cattle, using genomics and metabolomics data (Widmann *et al*, 2015). Moreover, our motif discovery analysis showed *TGFB1* co-expressed genes are mostly enriched for binding site of master regulators of muscle differentiation as *MEF2* and *MYOD*. Indeed, public available data show many of *TGFB1* target genes were associated with *MYOD* (Mullen *et al*, 2011). It is known signalling pathways are an effective mechanism for cells to respond to environmental cues by regulation gene expression. *TGFB1* signalling triggers the phosphorylation of *SMAD2/3* transcription factors, which co-bind with cell-type master regulators at the nuclear level allowing/triggering/leading to cell-type specific transcriptional changes (Schmierer & Hill, 2007; Mullen *et al*, 2011). In skeletal muscle cells, myoblasts and myotubes, *SMAD3* co-binds with *MYOD1* (Mullen *et al*, 2011). The overlap between *MYOD1* and *SMAD3* target genes demonstrate the significant association between both genes in skeletal muscle, in agreement with the tissue-specificity *TGFB1* signalling response (Mullen *et al*, 2011). The overlap percentage between our predicted binding sites and *MYOD1* Chip-seq data (18 and 24.5%) confirms previous analysis in mice where they reported only 20% of experimental validated distal enhancers in mouse myotubes with a bHLH (MyoD1) binding were actually bound by *MYOD* ChIP-seq data (Blum et al. 2012). Thus, suggesting additional transcription-factors and/or histone modification have a key role in *MYOD1* binding. The *SMAD3/MYOD1* co-bound regions for known target genes are also captured, such as the promoter regions of *ACTA1* and *ANKRD1*, both genes involved in skeletal muscle differentiation (Figure 4C). We also demonstrated predicted *MYOD1* binding regions are enriched for muscle regulatory regions across species (human, mouse and cow).

Altogether, we showed co-expressed genes with *TGFB1* are enriched for *SMAD3/MYOD1* binding sites, which we validate at the gene and enhancer level by proving not only *MYOD1* and *SMAD3* binding, but also their accessibility, in human, mouse and cow. In pigs, it has been indicated increased feed efficiency is associated with stimulation of muscle growth through TGF-β signalling pathway (Jing *et al*, 2015). Finally, although not directly co-expressed with *TGFB1*, oxytocin (*OXT*) was DE in muscle and despite the lack of knowledge on its role in this tissue, previous work in cattle have shown a massive increase of *OXT* expression in muscle of bovines chronically exposed to anabolic steroids (Jager *et al*, 2011). It is not known yet if oxytocin alone have an anabolic activity, but in a context where muscle growth seems to be associated with high FE animals, this is a hormone that worth further investigation.

From the 13 regulator genes that are DE between groups, six are involved in respiratory chain and are up-regulated in high FE group. Genes *ND1, ND4, ND4L, ND5, ND6* and *ND2,* which is DE but not identified as key regulator, are core subunits of the mitochondrial membrane respiratory chain Complex I (CI) which functions in the transfer of electrons from NADH to the respiratory chain, while *ATP8* is part of Complex V and produces ATP from ADP in the presence of the proton gradient across the membrane. Interestingly, greater quantity of mitochondrial CI protein were associated with high FE cattle by Ramos and Kerley (2013) whereas Davis et al. (2016) found higher CI-CII and CI-CIII concentration ratios for the same group. Other studies demonstrated high FE animals consume less oxygen (Chaves *et al*, 2015) and present lower plasma CO2 concentrations, which suggests a decreased oxidation process (Gonano *et al*, 2014). In general, the literature suggests mitochondrial ADP has greater control of oxidative phosphorylation in high FE individuals (Lancaster *et al*, 2014) and their increased mitochondrial function may contribute to feed efficiency (Connor *et al*, 2010). In pigs, differences in mitochondrial function were reported when analysing muscle (Vincent *et al*, 2015), blood (Liu *et al*, 2016) and adipose tissue transcriptomes (Louveau *et al*, 2016). Differences in metabolic rate associated with FE has long been discussed (Herd & Arthur, 2009) and here is corroborated by the up-regulation of *TSHB* in high FE animals, which stimulates production of T3 and T4 in thyroid thus increasing metabolism. It is inhibited by *SST*, a down-regulated hormone in this group which was also found to be differentially connected between high and low FE.

Looking at the DE genes, many hormones can be identified. Hormones are signalling proteins that are transported by the circulatory system to target distant organs in order to regulate physiology. Regarding the relationship between FE and other production traits of economic importance, *FSHB*, responsible for spermatozoa production by activating Sertoli cells in the testicles (Walker & Cheng, 2005), is up-regulated in low FE group and is inhibited by follistatin (*FST*), a gene found to be down-regulated in the same group. Moreover, in rats, it was already demonstrated *FSH* secretion is stimulated by somatostatin expression, which is up-regulated in low FE animals (Kitaoka *et al*, 1989). In this scenario, one could argue that selection for high FE delay reproduction traits, something that could be related to the lower fat deposition in this group, as previously observed (Alexandre *et al*, 2015; Santana *et al*, 2012; Gomes *et al*, 2012). Indeed, differences in body composition and in intermediary metabolism can impact on reproductive traits (Shaffer *et al*, 2011) and it has been observed before that feed efficient bulls present features of delayed sexual maturity, i.e. decreased progressive motility of the sperm and higher abundance of tail abnormalities (Montanholi *et al*, 2016; Fontoura *et al*, 2016). Moreover, high FE heifers presented less fat deposition and later sexual maturity which results in calving later in the calving season than their low FE counterparts (Randel & Welsh, 2013; Shaffer *et al*, 2011). It is important to point that low FE animals also present down-regulation of *AMH* and the fall of this hormone in serum was pointed as an excellent marker of Sertoli cells pubertal development (Rey *et al*, 1993).

Concerning the differences in lipid metabolism in divergent FE phenotypes, *FGF21*, a hormone up-regulated in liver of high FE animals, is associated in humans to decrease in body weight, blood triglycerides and LDL-cholesterol, with improvement in insulin sensitivity (Cheung & Deng, 2014). It is an hepatokine released to the bloodstream and an important regulator of lipid and glucose metabolism (Giralt *et al*, 2015). When we select its first neighbours in the network and perform an enrichment analysis we indeed found terms related to plasma lipoprotein particle remodelling, regulation of lipoprotein oxidation and cholesterol efflux mostly due to *FGF21* co-expression with the apolipoproteins *APOA4*, *APOC3* and *APOM*. In the same context, pro-melanin-concentrating hormone (*PMCH*) encodes three neuropeptides: neuropeptide-glycine-glutamic acid, neuropeptide-glutamic acid-isoleucine and melanin-concentrating hormone (*MCH*) being the last one the most extensively studied (Helgeson & Schmutz, 2008). *MCH* up-regulation has been related to obesity and insulin resistance, as well as increased appetite and reduced metabolism in murine models (Ludwig *et al*, 2001; Ito *et al*, 2003). *PMCH* gene is up-regulated in low FE animals and harbour SNPs found to be associated with higher carcass fat levels and marbling score (Walter *et al*, 2014; Helgeson & Schmutz, 2008).

In this work, we were able to identify several biological processes known to be related to feed efficiency, which together with the validation of the main transcription factors of the network, demonstrate the quality of the data and the robustness of the analyses, giving us the confidence to indicate candidate genes to be regulators or biomarkers of superior animals for this trait. The transcription factors NR2F6 and TGFB1 play central roles in liver and muscle, respectively, by regulating genes related to inflammatory response and muscle development and growth, two main biological mechanisms associated to feed efficiency. Likewise, hormones and other proteins secreted in plasma as oxytocin, adrenomedulin, TSH, somatostatin, follistatin and AMH are interesting molecules to be explored as potential biomarkers of feed efficiency.

## Material and methods

### Phenotypic data and biological sample collection

All animal protocols were approved by the Institutional Animal Care and Use Committee of Faculty of Food Engineering and Animal Sciences, University of São Paulo (FZEA-USP – protocol number 14.1.636.74.1). All procedures to collect phenotypes and biological samples were carried out at FZEA-USP, Pirassununga, State of São Paulo, Brazil. Ninety eight Nellore bulls (16 to 20 months old and 376 ± 29 kg BW) were evaluated in a feeding trial comprised of 21 days of adaptation to feedlot diet and place and a 70-day period of data collection. Total mixed ration was offered *ad libitum* and daily dry matter intake (DMI) was individually measured. Animals were weighted at the beginning, at the end and every 2 weeks during the experimental period. Feed efficiency was estimated by residual feed intake (RFI) which is the residual of the linear regression that estimates DMI based on average daily gain and mid-test metabolic body weight (Koch *et al*, 1963). Forty animals selected either as high feed efficiency (HFE) or low feed efficiency (LFE) groups were slaughtered on two days with a 6-day interval. Adrenal gland, hypothalamus, liver, muscle and pituitary samples were collected from each animal, rapidly frozen in liquid nitrogen and stored at −80 °C. Further information about management and phenotypic measures of the animals used in this study can be found in Alexandre et al. (2015).

### RNAseq data generation

Samples of nine animals from each feed efficiency group (high and low) were selected for RNAseq using RFI measure. For hypothalamus and pituitary, the nitrogen frozen tissue was macerated with crucible and pistil and stored in aliquots at −80 °C. Then, RNA was extracted using AllPrep DNA/RNA/Protein Mini kit (QIAGEN, Crawley, UK). For liver, muscle and adrenal gland, a cut was made in the frozen tissue and the RNA was extracted using RNeasy Mini Kit (QIAGEN, Crawley, UK). RNA quality and quantity were assessed using automated capillary gel electrophoresis on a Bioanalyzer 2100 with RNA 6000 Nano Labchips according to the manufacturer’s instructions (Agilent Technologies Ireland, Dublin, Ireland). Samples that presented an RNA integrity number (RIN) less than 8.0 were discarded.

RNA libraries were constructed using the TruSeq™ Stranded mRNA LT Sample Prep Protocol and sequenced on Illumina HiSeq 2500 equipment in a HiSeq Flow Cell v4 using HiSeq SBS Kit v4 (2×100pb). Liver, pituitary and hypothalamus were sequenced on the same run, each one in a different lane. Muscle and adrenal gland were sequenced in a second run, in different lanes.

### Gene expression estimation

The quality of the sequencing was evaluated using the software FastQC (http://www.bioinformatics.babraham.ac.uk/projects/fastqc/). Sequence alignment against the bovine reference genome (UMD3.1) was performed using STAR (Dobin *et al*, 2013), according to the standard parameters and including the annotation file (Ensembl release 89) and secondary alignments, duplicated reads and reads failing vendor quality checks were removed using Samtools (Li *et al*, 2009). Then, HTseq (Anders *et al*, 2014) was used to generate gene read counts and expression values were estimated by reads per kilobase of gene per million mapped reads (RPKM). Genes with average value lower than 0.2 FPKM across all samples and tissues were discarded.

Gene expression normalization was performed using the following mixed effect model (Reverter *et al*, 2005):

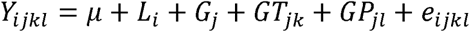

where the log2-transformed FPKM value for i-th library (86 levels), j-th gene (17,354 levels), k-th tissue (5 levels), l-th RFI phenotype (2 levels), corresponding to *Y_ijkl_*, was modelled as a function of the fixed effect of library (*L_i_*) and the random effects of gene (*G_j_*), gene by tissue (*GT_jk_*) and gene by RFI phenotype (*GP_jl_*). Random residual (*e_ijkl_*) was assumed to be independent and identically distributed. Variance component estimates and solutions to the model were obtained using VCE6 (Eildert Groeneveld, Milena Kovac and Norbert Mielenz, ftp://ftp.tzv.fal.de/pub/vce6/doc/vce6-manual-3.1-A4.pdf). Normalized mean expression (NME) values for each gene were defined as the linear combination of the solutions for random effects.

The mixed model used to normalize the expression data explained 96% of the variation in gene expression, of which the largest proportion (0.30) was due to tissue-specificity. Contrariwise, differences between high and low FE represented no variation (0.27E-11). For that reason, normalized mean expression (NME) were only used to identify tissue specific genes and the raw FPKM values were used for differential expression and co-expression analysis.

### Gene selection for network construction

In order to select a set of relevant genes for network analysis, we defined five categories based on the following inclusion criteria:

1. Differential expression (DE) - The mean expression value of each gene, for each group (high and low FE) and each tissue was calculated and then the expression of low FE group was subtracted from the expression in high FE group. Next, genes were ranked according to their mean expression in all samples for each tissue and divided in five bins. Genes were considered differentially expressed when the difference between the expression in high and low FE groups were greater than 3.1 or smaller than −3.1 standard deviation from the mean in each bin, corresponding to a t-test P<0.001.

2. Harbouring SNPs - Genes harbouring SNPs associated with feed efficiency, mainly indicated by GWAS, were identified using PubMed database (www.ncbi.nlm.nih.gov/pubmed/) and AnimalQTL database (www.animalgenome.org/cgi-bin/QTLdb/index) and only bovine data were considered regardless of breed.

3. Tissue specific (TS) - A gene was considered as tissue specific when the average NME in that tissue was greater than one standard deviation from the mean of all genes AND the average NME in all the other four tissues was smaller than zero.

4. Secreted - The human secretome database (www.proteinatlas.org/humanproteome/secretome) was used to select genes encoding proteins secreted in plasma by any of the analysed tissues (adrenal gland, hypothalamus, liver, muscle and pituitary).

5. Key regulators - In order to identify key regulatory genes to be included in the co-expression network, a list of genes were obtained from the Animal Transcription Factor Database (http://www.bioguo.org/AnimalTFDB/) and it was compared to a set of potential target genes in each tissue, composed by the categories: TS, DE, harbouring SNPs and secreted. The analysis was based on regulatory impact factor metrics (Reverter *et al*, 2010), which comprises a set of two metrics designed to assign scores to regulator genes consistently differentially co-expressed with target genes and to those with the most altered ability to predict the abundance of target genes. Those scores deviating ±1.96 standard deviation from the mean (corresponding to P<0.05) were considered significant. Genes presenting mean expression value less than the mean of all genes expressed were not considered in this analysis.

Some of the genes selected by the categories above were represented by more than one ensemble ID. Those duplications were removed for further analysis, keeping only the expression value of the most meaningful ensemble ID. Additionally, genes with mean expression across the samples equal to zero were also removed from further analysis.

### Co-expression network analysis

For gene network inference, genes selected by the five categories described previously were used as nodes and significant connections (edges) between them were identified using partial correlation and information theory (PCIT) algorithm (Reverter & Chan, 2008), considering all animals and all tissues. PCIT determinates the significance of the correlation between two nodes after accounting for all the other nodes in the network. The output of PCIT was visualized on Cytoscape (Shannon *et al*, 2003).

### Network validation through transcription factor biding motifs analysis

Using the regulatory impact factor metric (RIF) we prioritize key regulator genes from gene expression data and predict target genes based on co-expression network (PCIT). In order to assess whether those target genes were enriched for motifs associated to the top most connected regulators in the network with a DNA biding domain (transcription factors - TF), we performed motif discovery analysis in the set of co-expressed target genes (first neighbours of the TF) using i-cistarget method (Herrmann *et al*, 2012) and i-Regulon, a Cytoscape plug-in (Janky *et al*, 2014). These tools use human (hg19) as the reference species, therefore only genes with human orthology are assessed. Then, to validate the binding of the identified genomic regions by the TFs, we performed a region enrichment analysis across experimentally available TF bound regions from ChiP-seq in cell lines from the ENCODE consortium (1,394 TF binding site tracks). Finally, we converted identified enhancer regions to cow coordinates and searched for regions of open-chromatin using data from a public available studies in cow tissues.

### Differential connectivity

In order to explore differentially connected genes between high and low FE, two networks were created, one for each condition, using the same methodology described before. Then, the number of connections of each gene in each condition was computed and scaled so that connectivity varied from 0 to 1, making possible to compare the same gene in the two networks. The connectivity in high RFI group was subtracted from the connectivity in low RFI group and results deviating ±1.96 standard deviation from the mean were considered significant (P<0.05).

### Functional Enrichment

Functional enrichment analysis was performed on the online platform GOrilla (Gene Ontology enRIchment anaLysis and visuaLizAtion tool, http://cbl-gorilla.cs.technion.ac.il/), using all genes that passed FPKM filter as background, hypergeometric test and multiple test correction (FDR - false discovery rate). The human database was used to take advantage of a more comprehensive knowledge regarding gene functions. GO terms were considered significant when Padj<0.05. For genes in co-expression networks, visualized using Cytoscape (Shannon *et al*, 2003), the functional enrichment was performed with BiNGO plug-in (Maere *et al*, 2005) using the same background genes and statistical test.

## Acknowledgments

The authors thank Mrs. Elisângela C. M. Oliveira for all technical support. This study and PAA scholarships were funded by São Paulo Research Foundation (FAPESP) - Proc. 2014/07566-2, 2015/22276-3 and 2017/14707-0. M.N.S was funded by the CSIRO Science Excellence Research Office.

## Author contributions

HF was the overall project leader who conceived this study and supervised PAA in all data generation. TR performed the network analysis and supervised PAA in bioinformatics and data interpretation. MNS performed the motif discovery analysis. LRPN and JBSF provided informatics and statistical support. All authors contributed to and approved the final version of this manuscript.

## Availability of supporting data

Datasets supporting the results of this article is public available in the European Nucleotide Archive (ENA) as part of FAANG consortium under de study ID PRJEB27337 and can be accessed following the link https://www.ebi.ac.uk/ena/data/view/PRJEB27337.

## Supporting data

### S1 Supporting Information

**Information of reads mapping to bovine genome (UMD3.1) per sample.**

### S2 Supporting Information

**Scatter plots showing differentially expressed genes between high and low feed efficiency (FE) from adrenal gland, hypothalamus, liver, muscle and pituitary.** Dots represent the mean expression for a gene in high FE subtracted from the mean expression of the same gene in low FE (M) by the average expression value in both groups (A). Pink dots represent significant genes (P<0.001).

### S3 Supporting Information

**Differentially expressed genes between high and low feed efficiency in adrenal gland (A), hypothalamus (B), liver (C), muscle (D) and pituitary (E).**

### S4 Supporting Information

**Functional enrichment of the 248 genes down-regulated in high feed efficiency considering the five tissues (adrenal gland, hypothalamus, liver, muscle and pituitary).** Colour intensity increase with the significance of the term; white represents P>10^-3^ and the darkest orange represents P<10^-9^.

### S5 Supporting Information

**Genes selected for network construction for being differentially expressed between high and low feed efficiency (A), harbouring SNPs previourly associated with feed efficiency (B), tissue specific (C), coding proteins secreted in plasma (D) and key regulators (E).**

### S6 Supporting Information

**Functional enrichment for the 135 genes coding proteins secreted in plasma which the tissue of maximum expression is liver.** Colour intensity increase with the significance of the term; white represents P>10^-3^ and the darkest orange represents P<10^-9^.

### S7 Supporting Information

**Genes included in co-expression analysis.**

### S8 Supporting Information

**Top five key regulator genes network and enrichment. A) Network of genes *EPC1, NR2F6, MED21, ENSBTAG00000031687* and *CTBP1* and their first neighbours.** Nodes with diamond shape correspond to secreted proteins coding genes and triangles correspond to key regulators; all the other genes are represented by ellipses. Nodes with black borders are differentially expressed between high and low feed efficiency. Colours are relative to the tissue of maximum expression: blue represent liver, red represent muscle, yellow represent pituitary, green represent hypothalamus and orange represent adrenal gland. The size of the nodules is relative to the normalized mean expression values in all samples. Only correlations above 0.9 and bellow −0.9 and its respective genes are shown in this figure. **B) Functional enrichment of the 345 genes in the network (A).** Colour intensity increase with the significance of the term; white represents P>5×10^-3^.

### S9 Supporting Information

***TGFB1* network and enrichment. A) Network of *TGFB1* gene and its first neighbours.** Nodes with diamond shape correspond to secreted proteins coding genes and triangles correspond to key regulators; all the other genes are represented by ellipses. Nodes with black borders are differentially expressed between high and low feed efficiency. Colours are relative to the tissue of maximum expression: blue represent liver, red represent muscle, yellow represent pituitary, green represent hypothalamus and orange represent adrenal gland. The size of the nodules is relative to the normalized mean expression values in all samples. Only correlations above 0.9 and bellow −0.9 and its respective genes are shown in this figure. **B) Functional enrichment of the 157 genes in the network (A).** Colour intensity increase with the significance of the term; white represents P>5×10^-3^.

### S10 Supporting Information

***FGF21* network and enrichment. A) Network of *FGF21* gene and its first neighbours.** Nodes with diamond shape correspond to secreted proteins coding genes and triangles correspond to key regulators; all the other genes are represented by ellipses. Nodes with black borders are differentially expressed between high and low feed efficiency. Colours are relative to the tissue of maximum expression: blue represent liver, red represent muscle, yellow represent pituitary, green represent hypothalamus and orange represent adrenal gland. The size of the nodules is relative to the normalized mean expression values in all samples. Only correlations above 0.9 and bellow −0.9 and its respective genes are shown in this figure. **B) Functional enrichment of the 98 genes in the network (A).** Colour intensity increase with the significance of the term; white represents P>5×10^-3^.

### S11 Supporting Information

**NR2F6 i-cis Target results.**

### S12 Supporting Information

**NR2F6 predicted regions binding hg19.**

### S13 Supporting Information

**NR2F6 predcited regions bindg bosTau.**

### S14 Supporting Information

**TGBF1 i-cis Target results.**

### S15 Supporting Information

**MYOD predicted transcription factors binding sites in hg19.**

### S16 Supporting Information

**MYOD predicted transcription factors binding sites in mm8.**

### S17 Supporting Information

**MYOD predicted transcription factors binding sites in mm9.**

### S18 Supporting Information

**MYOD predicted transcription factors binding sites in bosTau6.**

### S19 Supporting Information

**Differentially connected genes between high and low feed efficiency.**

### S20 Supporting Information

**Functional enrichment for the 87 differentially co-expressed genes between high and low feed efficiency.** Colour intensity increase with the significance of the term; white represents P>10^-3^ and the darkest orange represents P<10^-9^.

